# An agent-based model of cattle grazing toxic Geyer’s larkspur

**DOI:** 10.1101/218032

**Authors:** Kevin E. Jablonski, Randall B. Boone, Paul J. Meiman

## Abstract

By killing cattle and otherwise complicating management, the many species of larkspur (*Delphinium* spp.) present a serious, intractable, and complex challenge to livestock grazing management in the western United States. Among the many obstacles to improving our understanding of cattle-larkspur dynamics has been the difficulty of testing different grazing management strategies in the field, as the risk of dead animals is too great. Agent-based models (ABMs) provide an effective method of testing alternate management strategies without risk to livestock. ABMs are especially useful for modeling complex systems such as livestock grazing management, and allow for realistic bottom-up encoding of cattle behavior. Here, we introduce a spatially-explicit, behavior-based ABM of cattle grazing in a pasture with a dangerous amount of Geyer’s larkspur (*D. geyeri*). This model tests the role of herd cohesion and stocking density in larkspur intake, finds that both are key drivers of larkspur-induced toxicosis, and indicates that alteration of these factors within realistic bounds can mitigate risk. Crucially, the model points to herd cohesion, which has received little attention in the discipline, as playing an important role in lethal acute toxicosis. As the first ABM to model grazing behavior at realistic scales, this study also demonstrates the tremendous potential of ABMs to illuminate grazing management dynamics, including fundamental aspects of livestock behavior amidst ecological heterogeneity.

## Introduction

The many species of larkspur (*Delphinium* spp. L.) present a serious, intractable, and complex challenge to livestock grazing management in the western United States [1–3]. Larkspur plants contain numerous norditerpinoid alkaloids, which are potent neuromuscular paralytics that, for reasons that are not entirely understood, are particularly effective at killing cattle, with yearly herd losses estimated at 2-5% for those grazing in larkspur habitat [3, 4]. To avoid such losses, producers will often abandon or delay grazing in pastures with larkspur, which creates a substantial opportunity cost and an impediment to achieving management objectives [1, 4].

Among the many challenges to improving our understanding of cattle-larkspur dynamics has been the difficulty of testing different grazing management strategies in the field. Not only is risking dead cattle impractical and unethical, but the complexity of livestock grazing management, especially when considered across the wide range of habitats and management regimes in which larkspur is found, suggests that results from individual field experiments would be unlikely to be broadly useful anyway [5, 6]. What is needed instead is a method of realistically testing grazing management strategies without risk to livestock and with the flexibility to test multiple scenarios. Agent-based models (ABMs) provide such a method.

ABMs are computational simulation tools that focus on the behavior of individual “agents” as they interact with one another and the environment [7]. They differ from other types of simulation models in being bottom-up (versus top-down) with group-level behaviors emerging from (usually) realistic individual behaviors rather than deterministic formulae [8]. ABMs are thus particularly useful in modeling complex systems, where the results of the interactions among system elements are not easily predicted or understood [9, 10]. Indeed, it has been suggested that bottom-up-simulation may be the best way to increase our understanding of complex systems, which is one of the most important challenges confronting modern science [9, 11, 12].

As noted by Dumont and Hill [11], ABMs are “particularly suited to simulate the behavior of groups of herbivores foraging within a heterogeneous environment”. The authors encourage the use of ABMs in situations where experimentation is impractical, and those where comparison of different management strategies is needed. Despite this encouragement, and despite the growing enthusiasm for ABMs in other disciplines, they have been little used in livestock grazing management research, despite the existence of relevant studies to parameterize such a model [e.g., 13–18]. This is at least partly due to confusion about the purpose and role of models in improving our understanding of complex systems.

Models can never be complete simulacra, and do not need to be in order to be useful. Instead, “models are neither true nor false but lie on a continuum of usefulness for which credibility can be built up only gradually” [19]. This credibility is built not just by model output but also, more importantly, through thoughtful model development. This ensures that the necessary simplification that occurs in modeling focuses in on rather than obscures the system processes of interest [20]. As noted by Augusiak et al. [19], in well-designed models the important question is the extent to which the model achieves its purpose in the light of existing evidence, rather than a binary yes or no regarding its validity.

Previous research into the relationship between grazing management and larkspur toxicosis has largely focused on timing of grazing, with some attention paid to mineral supplementation, pre-grazing with sheep, and, increasingly, genetic susceptibility [3, 4, 21–24]. Some papers have suggested that cattle behavior, influenced by management, can play a role in mitigating larkspur deaths [25, 26], but these ideas have received little empirical study. Only anecdotally has it been observed that, regardless of timing of grazing, it may be possible to eliminate losses to larkspur by increasing stocking density, due to a dilution effect (same amount of alkaloids, more cattle) or perhaps changes in herd behavior [27].

In this paper, we introduce a spatially-explicit, behavior-based ABM of cattle grazing in a pasture with a dangerous amount of Geyer’s larkspur (*Delphinium geyeri* Green), in which MSAL-type alkaloids are the dominant toxin [28, 29]. This model provides significant management-relevant insight for producers dealing with larkspur and demonstrates the great potential of ABMs to credibly model livestock grazing management dynamics, including fundamental aspects of livestock behavior amidst ecological heterogeneity.

## Methods

The model description follows the updated Overview, Design Concepts, and Details (ODD) protocol, an accepted method for standardizing published descriptions of ABMs [30].

### Purpose

We developed this model to test the effect of co-varying instantaneous stocking density [31] and herd cohesion (also known as troop length) [32] on cases of lethal acute alkaloid toxicosis caused by *D. geyeri*. Cases of lethal acute toxicosis are a product of intensity of exposure to alkaloids (via consumption) with passing time as a mitigating factor (via metabolism). Conceptually, this model functions as a mechanistic effect model (MEM) aimed at understanding the processes whereby toxic alkaloids kill grazing cattle. MEMs have been recognized for their potential to “close the gap between laboratory tests on individuals and ecological systems in real landscapes” [20]. We developed and executed the model in NetLogo 6.01, using the BehaviorSpace tool to implement simulations [33].

### Basic principles

Behavior-based encoding of cattle activities was the guiding principle of model design. As noted by Mclane et al. [7], “the behavior-based approach leads to a more complex web of decisions, and the responses of the animal to stimuli are often more multifaceted”. We add that the behavior-based approach is also more likely to allow for instructive emergent properties. In practice, the behavior-based approach means that at every step of the coding process we sought literature on actual cattle behavior and then encoded that behavior as realistically as possible. When literature was lacking we used our knowledge of cattle behavior from our years as livestock managers and researchers. The behavior-based approach also found expression in model evaluation, when one mode of evaluation was whether the cows in the model “act like cows”. This was achieved through a lengthy process of visual debugging and other implementation verification [19, 34].

A second core design principle was parsimony. Because this is the first ABM that we know of to incorporate cattle at the individual scale of interaction with the environment (1 m^2^) and extended to a realistic pasture size, we were initially tempted to include every cattle behavior we could. However, our focus on parsimony to the question at hand meant that we instead included only those behaviors relevant to the consumption of larkspur. A final guiding principle was that when a judgement call was needed, we erred on the side of making the effects of alkaloid toxicosis more prominent. If the model was to show an effect of grazing management on reducing larkspur-induced toxicosis, we wanted to be sure that we had taken every precaution against preconditioning it to do so.

Overall, we followed as closely as possible the process of “evaludation” laid out by Augusiak et al. [19], which is aimed at moving beyond insufficient and often counterproductive ideas about model validation to a more thorough process of generating credible models. Specifically, we incorporated data evaluation, conceptual model evaluation, implementation verification, output verification, and other analysis of model output.

### Entities and state variables

The model has two kinds of entities: pixels representing 1 m^2^ patches of land and agents representing 500 kg adult cows (1.1 animal-units). The patches create a model landscape that is 1663 x 1580 patches (1.66 km x 1.58 km, equal to 262.75 ha, of which 258.82 ha are within the pasture under study and 3.93 ha are outside the fence line and thus inaccessible). This landscape aims to replicate pasture 16 at the Colorado State University Research Foundation Maxwell Ranch, a working cattle ranch in the Laramie Foothills ecoregion of north-central Colorado that is a transition zone between the Rocky Mountains and the Great Plains. Several pastures on the ranch, including pasture 16, have significant populations of *D. geyeri*, which generate ongoing management challenges and have fatally poisoned cattle.

To make the model appropriately spatially explicit we included three sets of geographic data. First, using data from the Worldview-2 satellite (8-band multispectral, resolution 2 m) from July 10, 2016, we created an index of non-tree/shrub vegetation distribution within the pasture using a soil-adjusted vegetation index (SAVI) within ERDAS Imagine 2016 software at a resolution of 1 m [35, 36]. Second, as there are no developed watering locations in pasture 16, with ArcGIS Desktop 10.4 we digitized and rasterized (at 1 m) all locations of naturally occurring water as of July 2017 [37].

Lastly, in June and July of 2017 we mapped larkspur distribution and density in pasture 16 using a hybrid approach. We began by digitally dividing the pasture into 272 1-ha sampling plots. Because we knew larkspur to be of patchy distribution, in each plot we first mapped all larkspur patches (defined as areas with >1 larkspur plant • m^-2^) using an iPad equipped with Collector for ArcGIS 17.01 [38] and a Bad Elf Pro+ Bluetooth GPS receiver accurate to 2.5 m. To sample areas outside of larkspur patches for larkspur density, we counted all living larkspur plants in a 6-m-wide belt transect running horizontally across the plot, with the origin randomly assigned and any patches excluded [39]. Using ArcGIS Desktop we then extended the belt-transect-derived larkspur density to the rest of the plot (excluding patches), and both sets of data were integrated into a 1 m raster of larkspur distribution.

The number of cows (individual agents) in the model varies according to the chosen stocking density (SD, in AU • ha^-1^). Cows are assigned the role of “leader” (5%), “follower” (85%), or “independent” (10%) [16, 40, 41]. Each cow is also assigned a value for MSAL-tolerance and larkspur-attraction. MSAL-tolerance determines the MSAL-level at which a cow will “die” and is randomly assigned to create a normal distribution with 99.9% of values falling within 25% of the mean (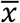 =4,000 mg, σ=333.33 mg) [42]. In this model, death does not result in the removal of a cow from the herd; instead, in order to preserve herd and other model functions it is recorded as having died, its MSAL-level is set to zero, and it continues to graze. Note that MSAL-tolerance can be understood as modeling genetic, physiological, and situational susceptibility.

Larkspur-attraction determines how much larkspur the individual cow will consume when in a patch with MSAL-content and is also randomly assigned to create a normal distribution with 99.9% of values falling within 25% of the mean (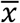=1.0, σ=0.083). A value of 1.0 means that the cow will consume larkspur at the same rate as other forage, while values greater or less than 1.0 cause the animal to, respectively, prefer or avoid larkspur. All functionally relevant state variables for patches and cows, as well as global variables and inputs, are described in Table 1.

**Table 1.** Relevant model variables.

| Entity | Variable | Description |
| --- | --- | --- |
| **Patches** | forage-mass | Amount of currently available forage (g) |
|  | n-forage-mass | Mean initial available forage in patches within a radius of 3 m (g) |
|  | MSAL-content | Amount of toxic alkaloids currently in patch (mg) |
|  | times-grazed | Number of times patch has been grazed |
| **Cows** | role | Role in the herd: leader, follower, or independent |
|  | MSAL-level | Current amount of MSAL alkaloids in cow's body (mg); metabolized with a half-life of one grazing-day |
| | MSAL-tolerance | Level at which cow will be recorded as having died (MSAL-level>MSAL-tolerance); assigned randomly from a normal distribution ($\bar{x}$=4,000 mg, $\sigma$=333.33 mg) |
| | larkspur-attraction | Factor determining the relative amount of larkspur a cow will eat when in a patch with MSAL-content; assigned randomly from a normal distribution ($\bar{x}$=1, $\sigma$=0.083) |
|  | herdmates | Agent-set consisting of nearest 20 cows |
|  | mean-herd-distance | Mean distance to herdmates |
|  | total-MSAL-intake | Total amount of MSAL alkaloids consumed during model run (mg) |
|  | daily-MSAL-intake | Amount of MSAL alkaloids consumed during current day (mg) |
|  | hydration | Hydration level, decreases to zero between visits to water |
|  | ready-to-go | Used by leader cows only, a measure of their inclination to move on from an overgrazed site |
| **Globals** | waterers | Patch-set of all watering locations |
|  | site-tolerance | Herd-size-dependent variable determining leader cows' tolerance for relatively overgrazed sites |
|  | site-radius | Radius of site when choosing a new site; product of herd-cohesion-factor and herd size resulting in space per cow ranging from 10 m<sup>2</sup> to 1000 m<sup>2</sup> |
|  | herd-distance | Desired mean-herd-distance; product of herd-cohesion-factor resulting in range from 10 m to 100 m |
| **Inputs** | kgs-per-hectare | Mean amount of usable forage (kg • ha<sup>-1</sup>) |
|  | mean-larkspur-mass | Mean mass of larkspur plants (g) |
|  | MSAL-concentration | MSAL alkaloid concentration in larkspur plants (mg • g<sup>-1</sup>) |
|  | herd-cohesion-factor (HCF) | Determines herd-distance and site-radius; range 1-10, increase leads to more cohesive herd |
|  | stocking-density (SD) | Instantaneous stocking density (AU • ha<sup>-1</sup>) |

### Scales

The model simulates cow activities at multiple temporal and spatial scales. In each tick (one cycle through the model code), each cow interacts with a single 1 m^2^ patch (a feeding station) by grazing (>99% of the time) or drinking water (twice per day) [13]. A tick does not represent time, but rather the occurrence of this interaction. This is because the duration of this interaction will vary depending on the amount of forage available, among other factors. Instead, time is represented by consumption of forage. When the average consumption of the grazing herd is equal to the average daily consumption of a 500 kg cow (12.5 kg), the model counts a grazing-day as having passed [43]. Total model run time is measured in animal-unit-months (AUMs) [44].

The narrowest scale of spatial interaction is the eating interaction occurring within a single patch (1 m^2^). When determining the next patch to graze, the cow’s decision is based on a desire either to move closer to its herdmates or to choose a nearby patch with maximum available forage. This decision happens on the scale of 2-25 m. Finally, leader cows make decisions on the scale of the entire pasture by deciding when it is time to visit water or time to move from the current feeding site to a new site.

Thus, there are four programmed spatial scales (additional scales may be emergent) at which the cows interact with the landscape: 1) the individual patch; 2) the scale of herd cohesion, set by the user; 3) the current feeding site; and 4) other feeding sites, identifiable by leader cows. The number of ticks that will pass before reaching a stopping point (say, 150 AUMs) depends on the number of animals grazing, their herd cohesion, the amount and distribution of available forage, and stochastic emergent properties of the model. For an expanded discussion of temporal and spatial scales of foraging behavior of large herbivores, see Bailey and Provenza [13].

### Process overview and scheduling

Fig 1 illustrates the model execution process for each tick. Each cow moves through each step of the process, but only performs those steps linked to its role.

**Fig 1.**
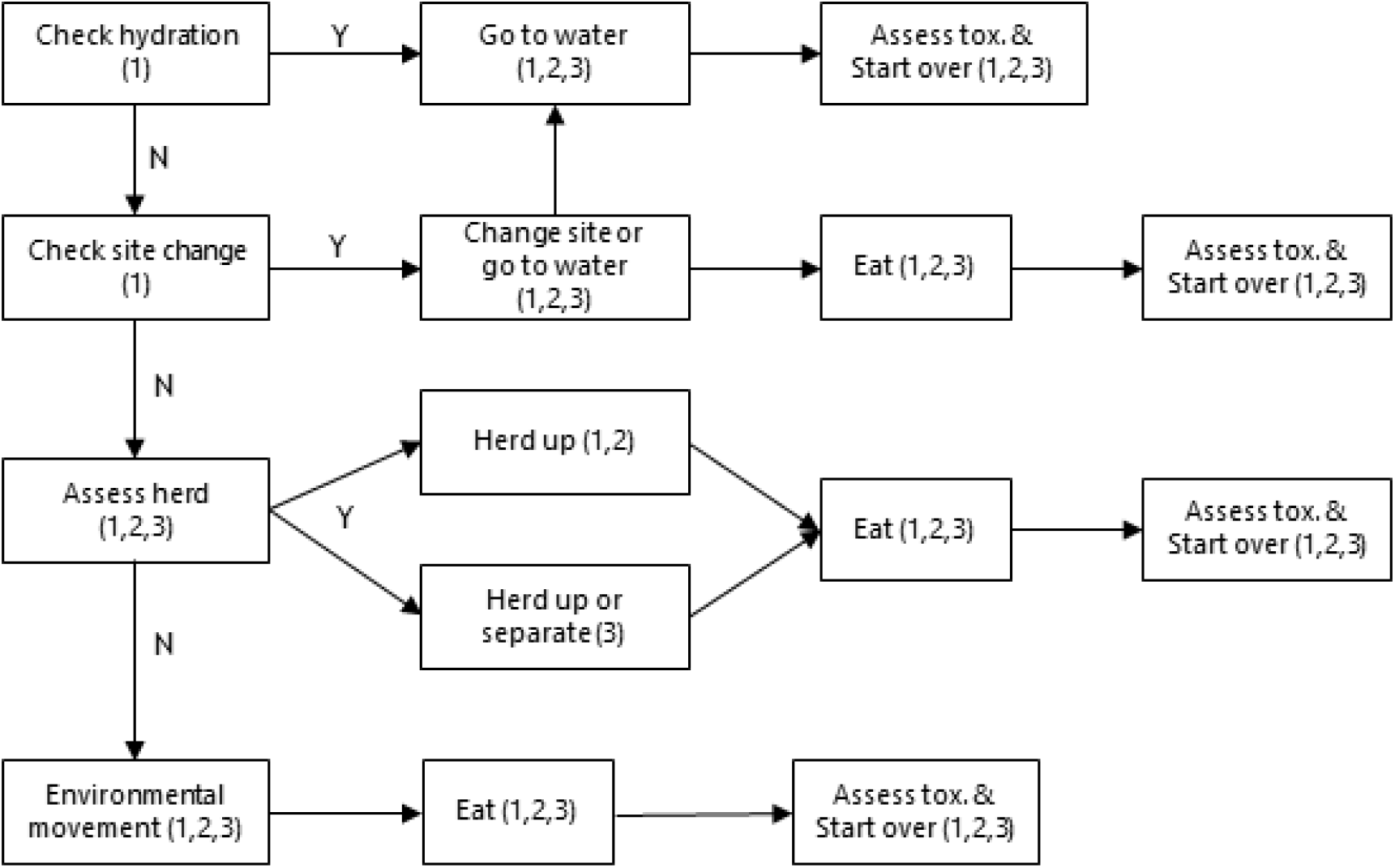
Pseudo-coded flow chart of model processes, with role of cows executing each process in parentheses. 1= leader, 2=follower, 3= independent.

### Check hydration

Each leader cow checks it hydration level, which is tied to forage consumption such that it depletes to zero twice per day. We chose two water visits per day based on personal communication about GPS collar data for the region [D. Augustine, USDA ARS, pers. communication; see 45]. If an individual leader detects its hydration level as less than or equal to zero, it initiates a movement to water for the whole herd.

### Go to water

The water source in pasture 16 is a stream that is intermittently below ground. The go-to-water procedure directs each cow to go to the nearest waterer patch with two or fewer cows already present. The hydration value for each cow is then set to maximum, and the value for ready-to-go for leader cows is set to site-tolerance – 1. This reflects the understanding that cattle will quickly graze and trample areas around water, rendering them unsuitable for grazing. Instead, they will pick desirable foraging areas in proximity to but not directly surrounding a watering site, expanding outward as these areas are grazed [13]. The model thus encourages a site change upon drinking water, but only if the area surrounding the watering site meets the criteria for increasing ready-to-go (explained below). A global variable ensures that no other processes occur during a tick when watering occurs.

### Check site change

This process is only executed by leader cows, each of which assesses the mean number of times patches within a radius of 10 m have been grazed. If these patches have been grazed relatively more (defined as >0.5 • mean times-grazed of all patches + 1.2) than the pasture as a whole, the value of ready-to-go increases by one. If this value reaches a pre-defined threshold (which increases with herd size), the individual then initiates a site change, but only if the individual’s hydration value is not approaching zero, in which case it instead initiates the go-to-water procedure. We arrived at the threshold formula for increasing the value of ready-to-go by using visual debugging and evaluation related to site change frequency, as well as theory on the optimization of grazing effort [13, 46].

If conditions for a site change are satisfied, the deciding leader cow first identifies the best five available sites, using criteria of number of times-grazed, forage-mass, and n-forage-mass to determine a centroid patch. The nearest of these patches is then used to create a new site at a radius that is linked to the user selected herd-cohesion-factor and the size of the herd, resulting in 10-1,000 m^2^ • cow^-1^ in the new site. The leader cow then initiates the change-site procedure for itself and all other cows.

### Change site

This procedure is initiated according to role, so that leader cows have first choice of their location in the new site, followers second, and independents third. Within the allocated new site, each cow chooses the patch with the most forage that has no cows on it or any of its four direct neighbors.

### Assess herd

In combination with the environmental-movement procedure, this process represents >99% of cow actions in the model. Each cow first sets its herdmates as the nearest 20 other cows [47]. For leader and follower cows, if the individual’s mean distance to these herdmates is greater than herd-distance, it “herds up”. This is achieved by facing the centroid of the herdmates and moving to the patch with maximum available forage that is 10-25 m in the direction of this centroid, within a cone of vision of ±45 degrees [14]. For independent cows, the same process occurs but is only initiated if the distance from herdmates is greater than 2.5 times the herd-distance of the other cows. Independent cows are also repelled from the center of their herdmates by moving away by the same procedure when they are within one-half of the herd-distance.

### Environmental movement

If none of the above procedures are implemented, each cow will make a movement decision based on local grazing conditions. If the patches within a radius of 10 m are relatively ungrazed (mean times-grazed < 0.5) the cow will move to the patch with the most available forage within 2 m, within a ±45 degree cone of vision [13]. If the same area is relatively well grazed (mean times-grazed ≥ 0.5), the cow then looks further afield, choosing the patch with the most available forage within 10 m, within a cone of vision of ±45 degrees.

### Eat

The eat procedure is the core interaction between the cows and the forage, both non-larkspur and larkspur. Behavior varies slightly depending on how many times the patch has previously been grazed. If the current visit is the first time it has been grazed, the cow eats 40% of the available forage [15, 18]. If it is the second visit, it eats 50% of what remains. In the third and any subsequent visits, it eats 60%. Each cow then increases its consumption-level by the same amount and decreases its hydration value. If there is larkspur present (in the form of MSAL-content), that is consumed according to the individual cow’s larkspur-attraction value, increasing the MSAL-level of the cow. The corresponding patch values are decreased to account for consumption. Lastly, times-grazed in the patch is increased by one.

### Assess toxicosis

This process is triggered at the end of each grazing-day for all cows in order to assess their toxicosis status, which is measured as their MSAL-level relative to their MSAL-tolerance. Note that MSAL-level is measured continuously throughout the model run, and has an elimination half-life of one grazing-day [48]. If MSAL-level exceeds MSAL-tolerance, the count of deaths for the model run is increased by one, MSAL-level is set to zero, and the cow continues. Numerous other data on toxicosis status are recorded for all cows at this point. Lastly, the MSAL-level for each cow is multiplied by 0.5.

## Design concepts

### Emergence

Because the actions of the cows are encoded via simple behavior-based processes, nearly all model patterns can be considered emergent. These include the stochastic distribution of the herd and subherds, forage consumption, larkspur consumption, grazing pressure and patterns, and site changes. Assessment of these un-coded emergent properties and patterns was critical to establishing the credibility of the model [20].

### Adaptation, objectives, learning, and predictions

The cows adapt to the grazing environment as they and their fellow cows graze, continually seeking their main model objective of maximizing forage consumption within behavioral limits [14]. There is no encoded learning or prediction, as the cows are programmed to be familiar with the location of forage and water in the pasture. However, it may be that learning and prediction are emergent, in that activities that we might consider to be evidence of those behaviors are visible in the model as a result of the simple encoded behaviors.

### Sensing and interaction

The cows sense each other and their environment at multiple spatial scales. Interaction occurs with other cows whenever moving to a new patch, both via sensing if a patch is already occupied and by seeking to herd up when too far from their herdmates.

### Stochasticity

There is no environmental stochasticity in this model iteration, as we sought to make the landscape as realistic as possible by incorporating relevant data from the real pasture 16. However, cattle interactions with the forage and larkspur demonstrate moderate stochasticity.

### Initialization

Landscape initialization begins by loading the SAVI layer and a user-input value for available forage per ha (kgs-per-hectare). The model uses a nonlinear exponential formula to distribute forage such that the patches with the least forage contain one-third of the mean forage, while the patches with the most contain three times the mean forage. Next, the model incorporates the larkspur distribution layer, using inputs of median larkspur mass (g) and mean MSAL concentration (mg • g^-1^) to generate an MSAL alkaloid (hereafter simply “alkaloid”) content for each patch. These values are based on our unpublished data on *D. geyeri* mass and toxicity at the Maxwell Ranch such that larkspur plants in areas of high SAVI were 50% larger than the median, and larkspur plants in areas of low SAVI were 50% smaller than the median. Finally, the model incorporates the water location layer. All other patch variables are derived from these inputs. Fig 2 shows the initialized landscape.

**Fig 2.**
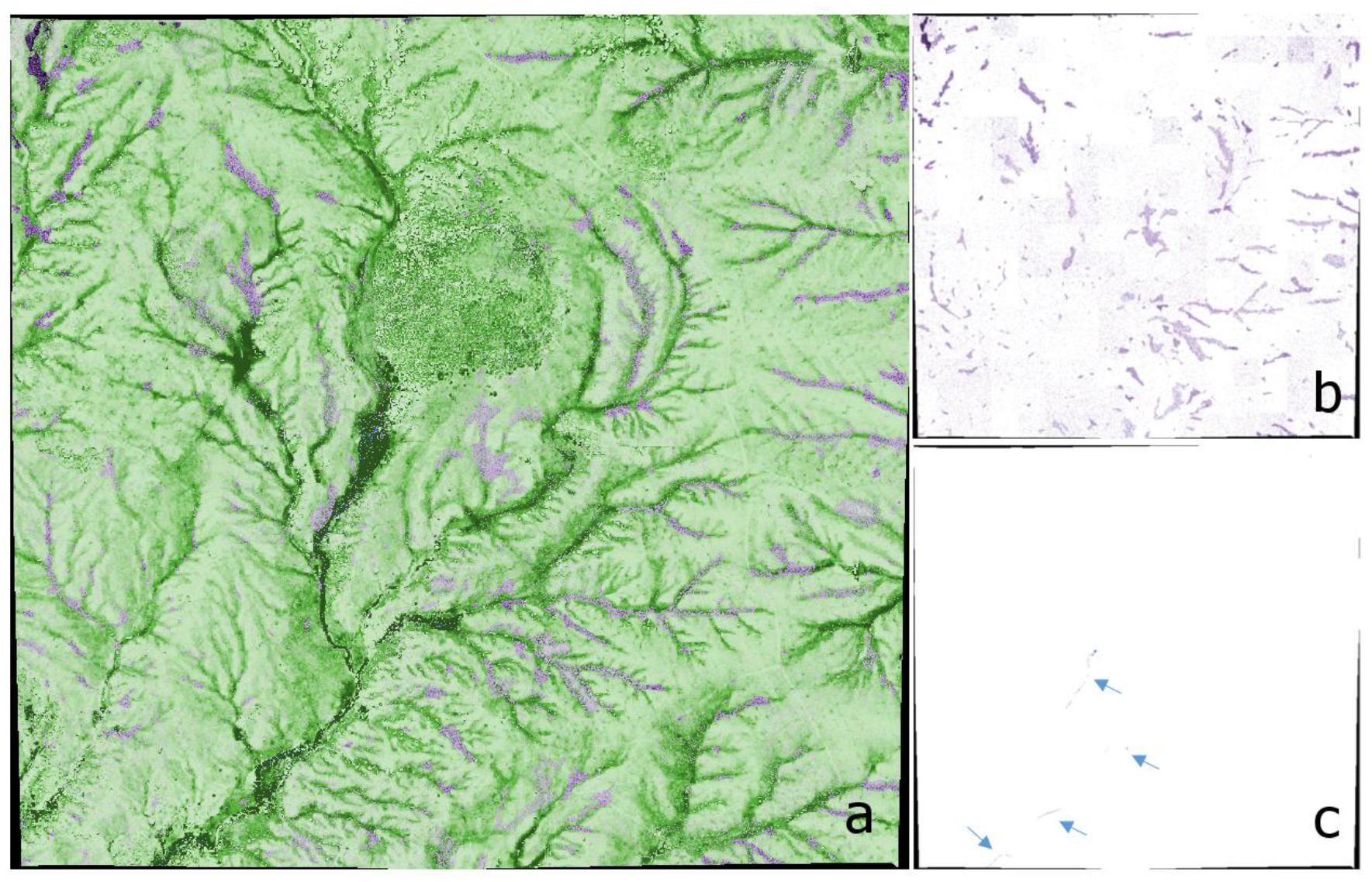
Model landscape, 1.66 km x 1.58 km. (a) Initialized full model landscape, with darker green indicating areas with greater aboveground forage biomass. (b) Landscape with larkspur locations only, with darker purple representing higher MSAL-content and with results of hybrid sampling method evident. (c) Landscape with watering locations only, pointed out by arrows.

The final step in model initialization is to create the cows by using the input of stocking-density multiplied by the area of the pasture. All cows are initially in the same random location in the pasture. This location is largely irrelevant as the cows immediately go to water, but we did not want it to be the same location each time because this would be unrealistic (pasture 16 has multiple entrances for cattle) and would limit stochasticity. At this point, the model is fully initialized and is executed following the processes laid out above.

### Simulation

We used the BehaviorSpace tool in NetLogo to run a full factorial simulation of four different levels of both herd-cohesion-factor (1, 4, 7, and 10) and stocking-density (0.25, 0.5, 1.0, and 2.0 AU • ha^-1^). We replicated each combination 30 times, for a total of 480 simulations. Input median larkspur mass was 3.5 g and input MSAL alkaloid concentration was 3.0 mg • g^-1^. We chose these values to be representative of an excellent growing year with larkspur plants at bud stage, when the alkaloid pool (total available mg) is highest—arguably the most dangerous possible conditions. This is also a time of year that cattle grazing in larkspur habitat is frequently avoided, despite being a highly desirable time for grazing [1, 4, 49]. Input value for kgs-per-hectare was 500 kg, based on current ranch usage and typical values for the area.

### Observation

Of primary importance were data related to alkaloid consumption, assessed according to dose-response data from previous research [42]. Most interesting was the number of times in a model run that any individual cow crossed the threshold into potentially lethal acute toxicosis, during which they would be expected to be recumbent and unable to stand, with a high likelihood of death [42]. To measure the number of such cases, the model counted cows whose MSAL-level exceeded their MSAL-tolerance at the end of a grazing-day.

The model also recorded data underlying the trends found for lethal acute toxicosis, most importantly data on daily, total, and maximum alkaloid intake. These data assisted in identifying potential mechanisms for the role of herd cohesion and stocking density in influencing deaths. Additional data, such as forage consumption, number of site changes, travel distance per day, and evenness of grazing impact, provide additional insight and model output verification.

### Statistical analysis

We used both JMP Pro 13.0.0 and R statistical software, version 3.3.3 for data analysis and presentation [50, 51]. Data for daily alkaloid intake, which amounted to 1.88 million data points, were organized and cleaned using OpenRefine 2.8 [52]. We began by assessing the role and relative influence of HCF and SD in generating lethal acute toxicosis, within two contexts: first, using their 16 combinations as “management levels” to explore overall trends in a management-relevant manner; and, second, using HCF and SD as continuous variables within a regression framework to provide more information on the relative influence of each. To regress the lethal acute toxicosis count data we used a generalized linear model (GLM) with a negative binomial distribution and a log-link function using the MASS package in R [53, 54]. To confirm that the negative binomial distribution was the correct choice, we compared it to a GLM with a Poisson distribution and a log-link function. The GLM with the negative binomial distribution was superior, using residual deviance and Aikaike’s information criterion (AIC) as judgment criteria [55].

To identify mechanisms for how HCF and SD were influencing deaths, we used the same negative binomial GLM approach to analyze the relationship between various intake data and lethal acute toxicosis. We did so by first hypothesizing which factors were driving deaths, and then looked at single-factor models for each, assessed using AIC values and model coefficients [55]. Because the goal was to identify key mechanisms rather than determine the best predictive model, this provided more insight than examining a global model or various permutations of factors.

Finally, we analyzed the relationship of HFC and SD to the identified mechanisms using multiple linear regression (R base package). While there were some indications of heteroscedasticity and outliers, we determined that linear regression was robust to those errors in these cases. We confirmed this by also fitting alternate models within other regression frameworks (robust and non-parametric), which returned very similar results. Interaction effects are shown when significant; otherwise, they were excluded from the models.

## Results

### Model output verification

A core element in the evaluation of behavior-based mechanistic effect models is a comparison between multiple emergent model patterns and observed patterns in the real system [20]. In this case, this helped to establish that the modeled cows, coded for individual behaviors, acted like real cows when interacting with one another and the landscape, at least in regard to behaviors relevant to larkspur consumption. Toward this end, first we offer Fig 3 to illustrate how varying HCF influences herding patterns, and to show how grazing was distributed across the pasture in one model run.

**Fig 3.**
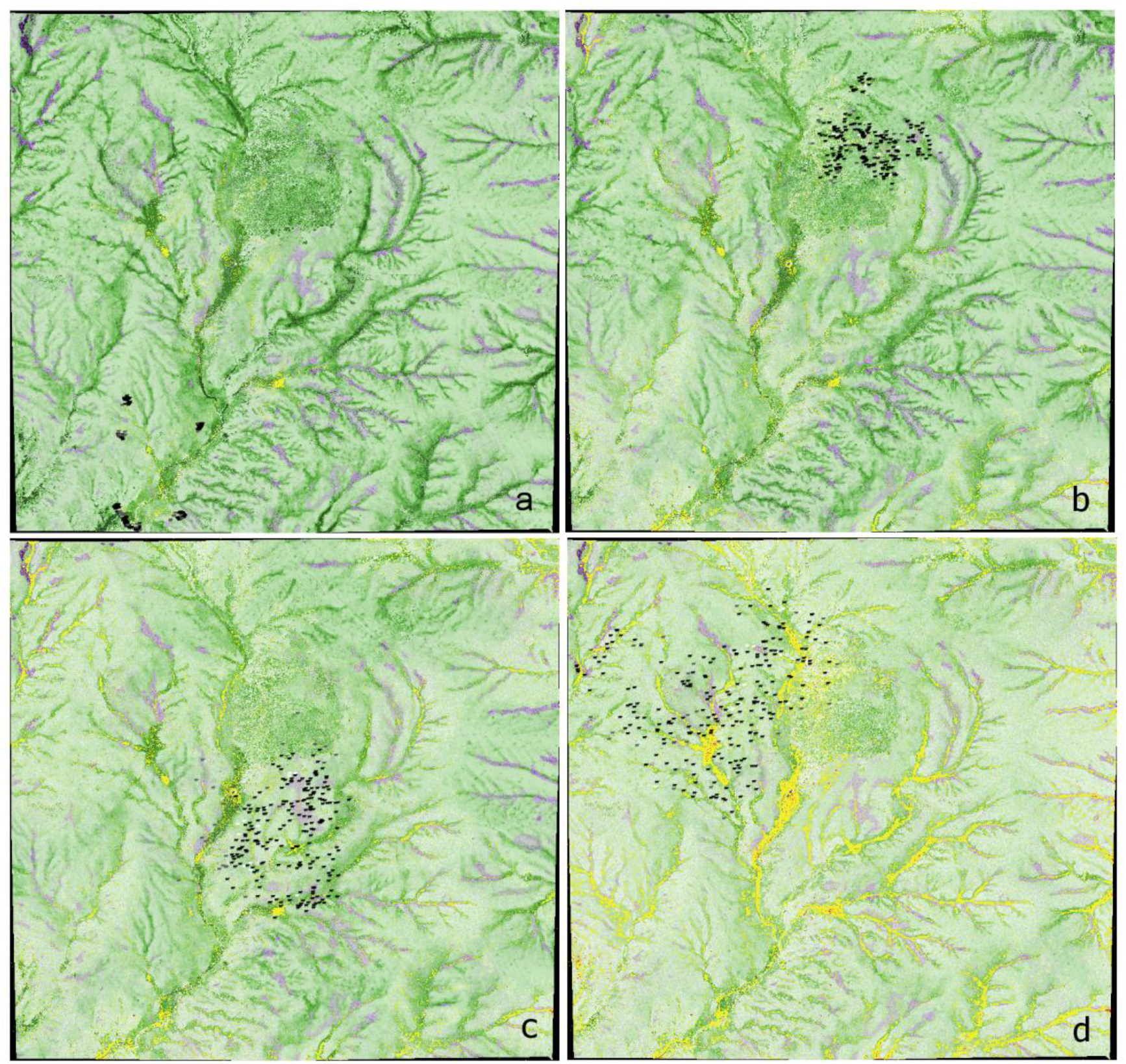
The effect of varying herd-cohesion-factor (HCF) on herd patterns, displayed at different levels of pasture usage (AUMs). Note that the cows depicted in these images are drawn 200 times larger than they really are to aid visualization, which makes them appear closer to one another than they are. Pasture size is 1.66 km x 1.58 km, and stocking density for all images is 1.0 AU • ha^-1^. White cows are leaders, black followers, and gray independents. Yellow indicates patches that have been grazed twice, red three times. (a) HCF=10, AUMs=14; (b) HCF=7, AUMs=68; (c) HCF=4, AUMs=119; (d) HCF=1, AUMs=163. Typical usage for this pasture (258.82 ha) is 150 AUMs.

Decreasing HCF increases overall herd separation and leads to more wandering among the independent cows and others. Note that in Fig 3a the cows have formed distinct subherds. This appears to be an emergent property of cows grazing with high herd cohesion (herd-distance ≤ 20 m).

The cows initially graze the areas with high forage amounts (dark green) in relative proximity to the water, and gradually extend their impact outward, targeting high productivity areas. By the end of the grazing cycle (Fig 3d), they have visited the entire pasture, though areas furthest from water have been grazed less [56]. Areas of initial high forage mass have been grazed two or more times, while many areas of low forage mass have not been grazed at all. These results are in line with well-established qualitative understanding of grazing patterns in large pastures [13, 44].

The variation in forage consumption among individuals also aligned well with the variation seen in real cows foraging native pasture. While a grazing-day for the whole herd was defined as mean consumption of 2.5% of body weight (12.5 kg), the mean 99.9% daily range of consumption for all model runs was 2.34-2.66% of body weight. This range of consumption aligns well with common “rules of thumb” and predictive formulae [43, 57, 58].

The mean value for site changes per day for the 16 management levels varies from 2.3 for few cows grazing very loosely (HCF=1, SD=0.25) to 6.0 for many cows grazing very cohesively (HCF=10, SD=2.0). These values are in line with the estimate of 1-4 hours per feeding site by Bailey and Provenza [13]. For runs with few cows grazing with little cohesion (HCF=1, SD=0.25), mean daily travel was 4.16 km, while many cows grazing very cohesively (HCF=10, SD=2.0) traveled an average of 7.40 km per day. These numbers and the positive trend also track well with data from previous studies [59].

As a last point of output verification, we were interested to see if the number of modeled cases of larkspur-induced lethal acute toxicosis would parallel numbers from the literature when we modeled grazing to be similar to the current management scheme. When modeled to reflect current management practices, with HCF=4, SD=0.5, and for 150 AUMs (removing approximately 45% of available forage), we recorded a mean of 2.8 cases of lethal acute toxicosis across 30 model iterations. This amounts to 2.4% of cows, which falls within the estimate of 2-5% in pastures with dangerous amounts of larkspur [4]. Additionally, individual model runs of zero deaths occurred in all but four of the management levels, which aligns with our anecdotal understanding of producer experience.

### Lethal acute toxicosis

On its own, increased herd cohesion demonstrated the potential to significantly reduce deaths. For example, at a stocking density of 0.5 AU • ha^-1^, mean deaths declined from 4.33 at HCF=1 to 1.37 at HCF=10. Similarly, increased stocking density in the absence of changes in herd cohesion also greatly reduced deaths, for example from a mean of 7.5 at SD=0.25 to 0.70 at SD=2 at a constant HCF of 4. Working together, increases in both herd cohesion and stocking density from the minimum to the maximum achieved a 99.6% reduction in deaths (Fig 4). The mean value for MSAL-tolerance among dead cows was 3,725.8 mg, while the mean value for larkspur-attraction was a factor of 1.06. Of 1,132 total deaths in the simulation, 3.9% were among cows with the role of leader, 78.7% were among followers, and 17.4% were among independents.

**Fig 4.**
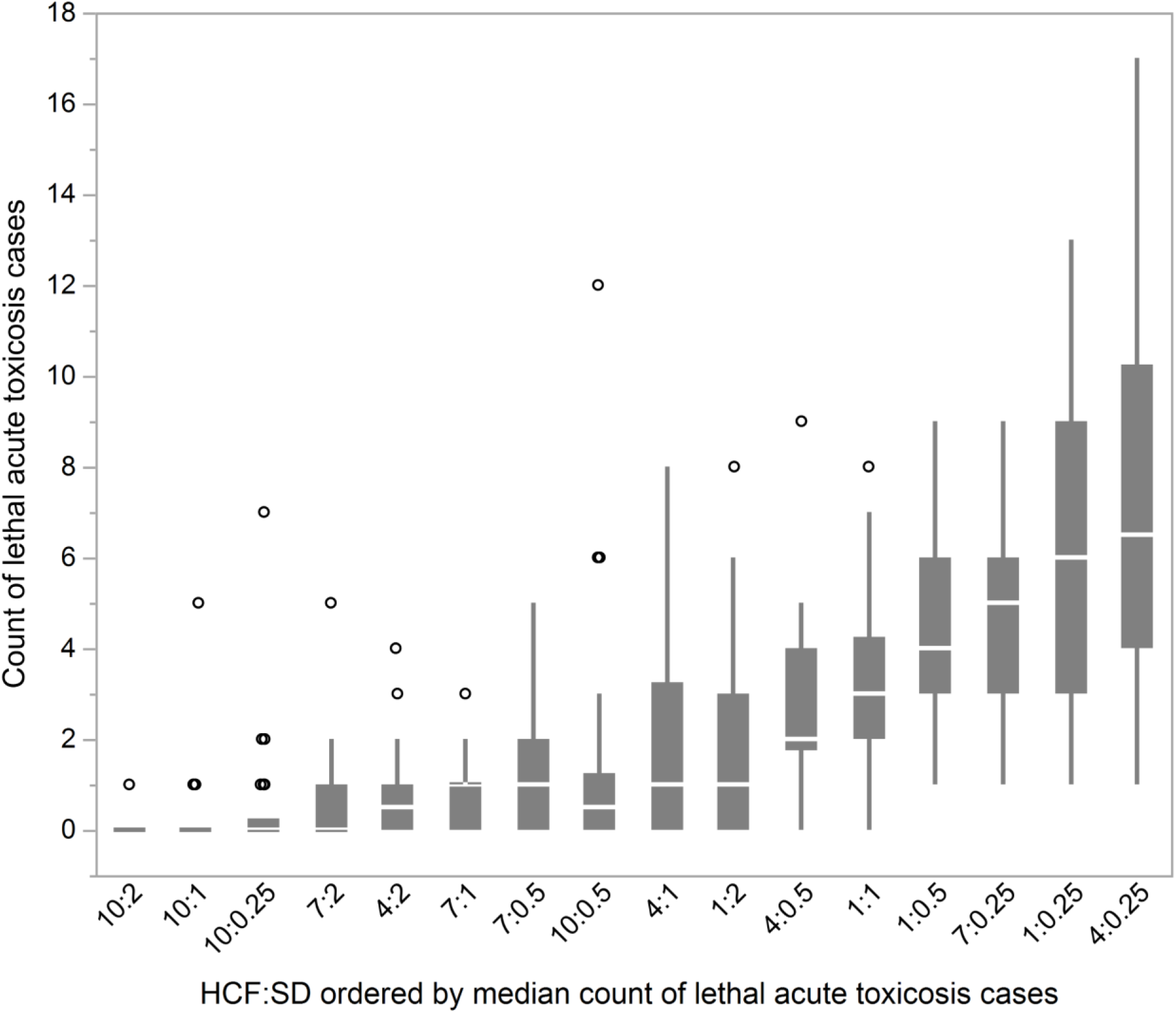
Box plots of distribution of counts of lethal acute toxicosis cases (MSAL-level ≥ MSAL-tolerance at end of grazing-day). From 30 model runs for each combination of herd-cohesion factor (HCF) and stocking-density (SD), ordered by median count of lethal acute toxicosis cases, with outliers as jittered circles.

The coefficient for HCF (Table 2), as a log odds ratio, indicates that an increase of one in HCF resulted in a 13.5% decrease in occurrences of lethal acute toxicosis. The coefficient for SD indicates that an increase of one in SD resulted in a 54.8% decrease. Lastly, the coefficient for the interaction of HCF with SD indicates that an increase in either HCF or SD slightly increases the effect of the other. The GLM β coefficients indicate that HCF had 91.8% of the influence of SD in reducing deaths.

**Table 2.** Results of GLM with negative binomial distribution and log-link function for count of lethal acute toxicosis as predicted by herd-cohesion-factor (HCF) and stocking-density (SD). β coefficients are from the same GLM without the interaction present. GLM fit: Fisher scoring iterations=1; residual deviance=516.94 on 476 degrees of freedom; AIC=1686.3.

| Coefficient | Estimate | Std. error | p-value | $\beta$ |
| --- | --- | --- | --- | --- |
| Intercept | 2.341 | 0.128 | <0.001 |  |
| HCF | -0.145 | 0.024 | <0.001 | -0.225 |
| SD | -0.793 | 0.136 | <0.001 | -0.245 |
| HCF:SD | -0.079 | 0.029 | 0.007 |  |

### Identifying mechanisms

We hypothesized that five factors might explain how HCF and SD were reducing deaths: mean individual daily alkaloid intake (the average single-day alkaloid intake in a model run), standard deviation of individual daily alkaloid intake, mean maximum individual daily alkaloid intake (each cow’s worst day), standard deviation of maximum individual daily alkaloid intake, and the coefficient of variation for individual total alkaloid intake. Results for the comparison of single-factor models reveal varying influence on lethal acute toxicosis among these factors (Table 3).

**Table 3.**
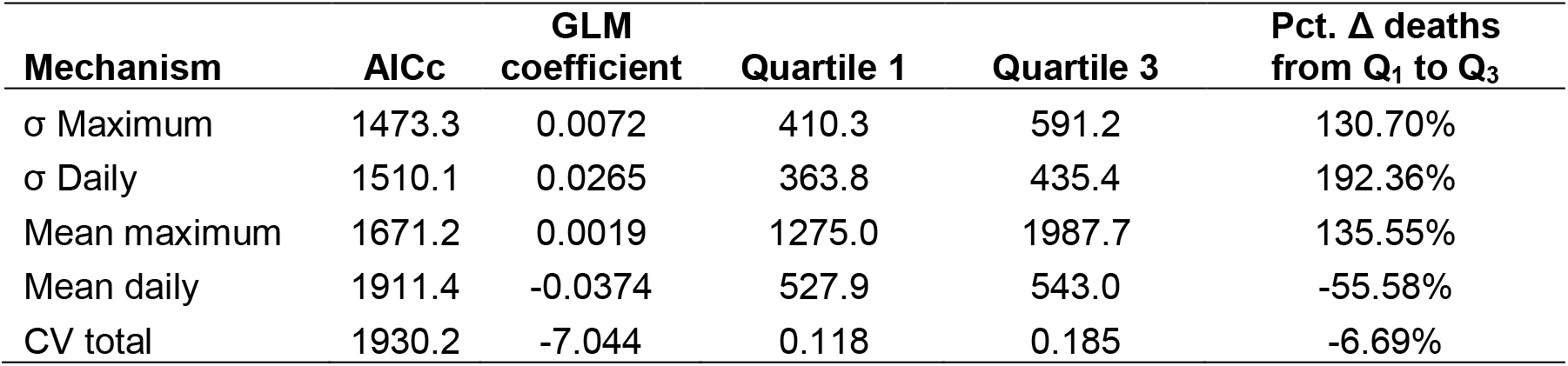
Results for comparison of single-factor negative binomial generalized linear models with a log-link function using corrected Aikaike’s information criterion. All values for quartiles are in mg, except for CV total, which is unitless. Percent Δ deaths from Q_1_ to Q_3_ is observed percent change in lethal acute toxicosis count between quartile one and three.

Because they had the most significant effect on lethal acute toxicosis, and were scored lowest for AICc, we focused the rest of the analysis on examining the relationship of HCF and SD to standard deviation of maximum individual daily alkaloid consumption, standard deviation of individual daily alkaloid consumption, and mean maximum individual daily alkaloid consumption. A model for lethal acute toxicosis count that contained these three mechanisms had an AICc score of 1368.3.

### Daily alkaloid intake

Mean individual daily alkaloid intake represents the mean of every single-day alkaloid intake for every cow, and ranged from a low of 525.1 mg (HCF=4, SD=0.25) to a high of 550.9 mg (HCF=10, SD=0.25). Multiple linear regression results indicate that HCF and SD had limited influence on mean daily intake (adj. R^2^<0.19), with both associated with slight increases. On the other hand, the standard deviation of daily alkaloid intake, which quantifies the spread of the distribution of daily alkaloid intake values, differed significantly between management levels, from a high mean of 460.5 mg (HCF=1, SD=0.25) to a low mean of 301.3 mg (HCF=10, SD=2). Multiple linear regression results indicate that HCF and SD were strongly influential, with HCF exerting 93.0% more influence than SD (Table 4).

**Table 4.** Results of multiple linear regression for the standard deviation of individual daily alkaloid intake as predicted by herd-cohesion-factor (HCF) and stocking-density (SD). Adj. R^2^=0.76.

| Coefficient | Estimate | Std. error | p-value | $\beta$ |
| --- | --- | --- | --- | --- |
| Intercept | 487.79 | 2.61 | <0.001 |  |
| HCF | -11.33 | 0.33 | <0.001 | -0.774 |
| SD | -29.38 | 1.64 | <0.001 | -0.401 |

A box plot showing the distribution of all individual daily alkaloid intake values (*n=1.88* • 10^5^) at each management level further illustrates these patterns (Fig 5).

**Fig 5.**
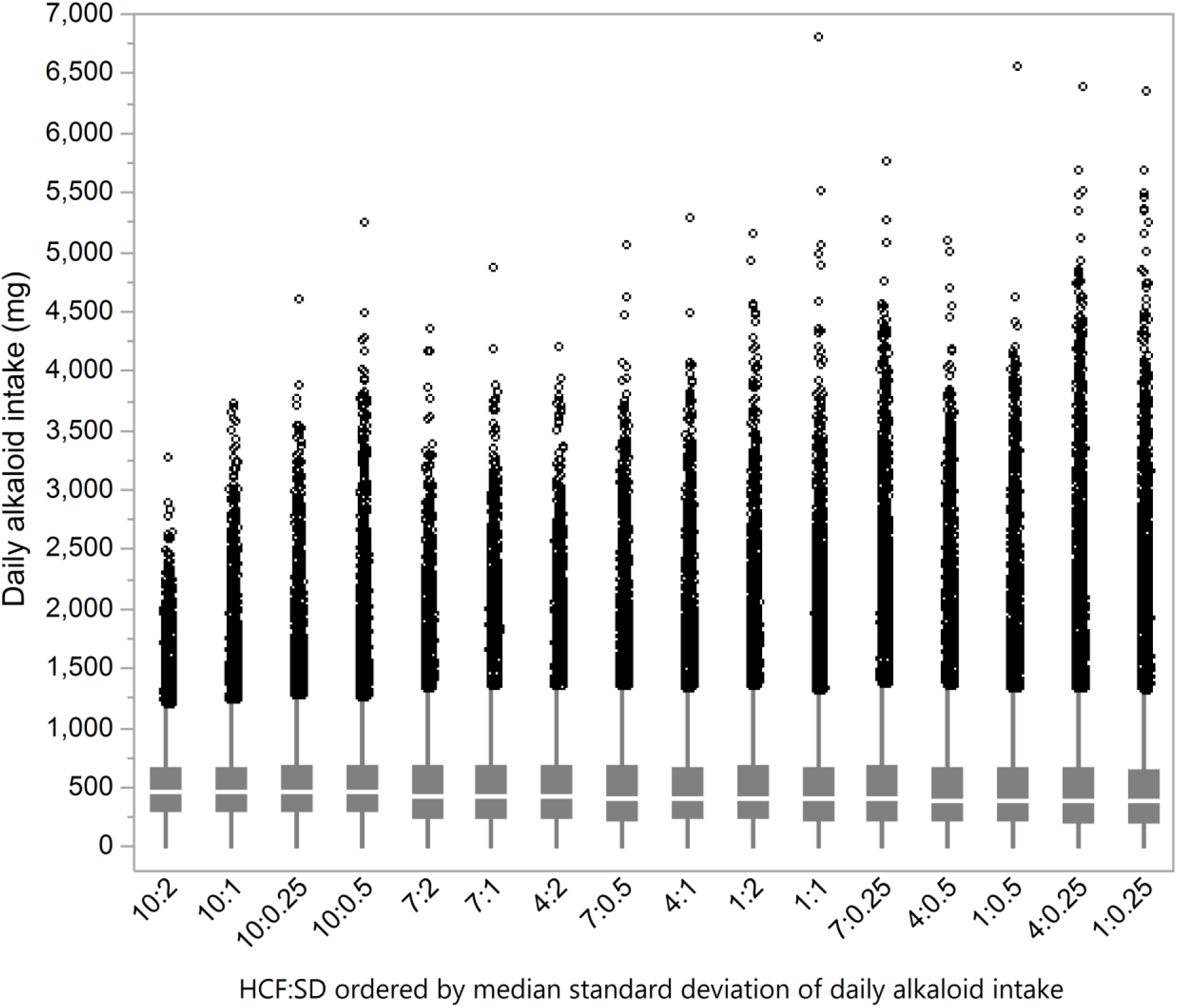
Box plots of distribution of individual daily alkaloid intake (mg; *n*=1.88 • 10^-5^). From 30 model runs for each combination of herd-cohesion factor (HCF) and stocking-density (SD), ordered by median standard deviation of daily alkaloid intake, with outliers as jittered circles.

### Maximum daily alkaloid intake

Mean maximum individual daily alkaloid intake quantifies the mean worst day for all cows during a model run, and ranged from 1,045.6 mg (HCF=10, SD=2) to 2,450.2 mg (HCF=1, SD=0.25). The standard deviation of maximum individual daily alkaloid intake quantifies how widely dispersed this value was among the herd members, and ranged from 303.0 mg (HCF=10, SD=2) to 704.0 mg (HCF=4, SD=0.25). Regression results for both factors provide further insight into the relationship of HCF and SD to lethal acute toxicosis (Tables 5–6).

**Table 5.** Results of multiple linear regression for the mean of maximum individual daily alkaloid intake as predicted by herd-cohesion-factor (HCF) and stocking-density (SD). Adj. R^2^=0.82. β coefficients are from the same model without the interaction present.

| Coefficient | Estimate | Std. error | p-value | $\beta$ |
| --- | --- | --- | --- | --- |
| Intercept | 2547.56 | 28.58 | <0.001 |  |
| HCF | -61.86 | 4.44 | <0.001 | -0.31 |
| SD | -686.15 | 24.80 | <0.001 | -0.84 |
| HCF:SD | 22.75 | 3.85 | <0.001 |  |

**Table 6.** Results of multiple linear regression for the standard deviation of maximum individual daily alkaloid intake as predicted by herd-cohesion-factor (HCF) and stocking-density (SD). Adj. R^2^=0.47. No significant interaction was present.

| Coefficient | Estimate | Std. error | p-value | $\beta$ |
| --- | --- | --- | --- | --- |
| Intercept | 718.57 | 11.33 | <0.001 |  |
| HCF | -22.34 | 1.42 | <0.001 | -0.52 |
| SD | -96.06 | 7.12 | <0.001 | -0.45 |

### Distinct persistent subherds

Model outputs (Fig 6) suggested an apparent scalar behavioral discontinuity between HCF=7 and HCF=10, which we believe results from the emergent property of distinct persistent subherds.

**Fig 6.**
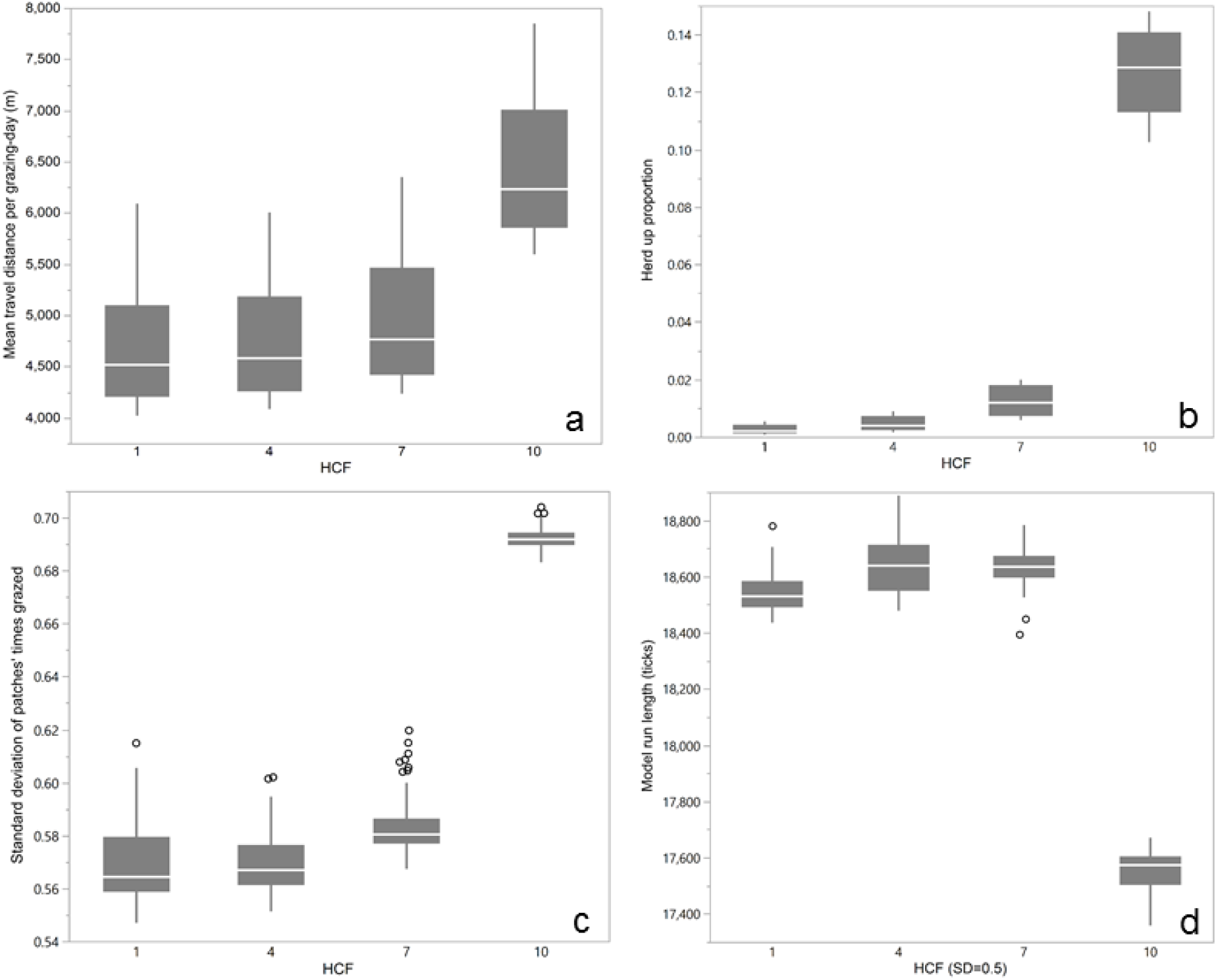
Box plots of various model evaluation data demonstrating effect of distinct persistent subherds. (a) Mean individual travel distance per grazing day (m) by herd cohesion factor (HCF); (b) Proportion of use of assess herd procedure (versus environmental movement) to choose a new grazing patch, a measure of herd-based versus individual optimization, by HCF; (c) Standard deviation of times-grazed count for all patches at end of model run, a measure of grazing heterogeneity, by HCF; (d) Total model run length, an inverse indicator of grazing efficiency, by HCF at stocking density=0.5.

## Discussion

Research into best practices for grazing management in larkspur habitat has long focused on either attempts to eliminate larkspur or on phenological avoidance (what we term “fight or flight”). Because elimination through herbicides or mowing is costly and often impractical [60], most research and current recommendations focus on avoiding grazing in larkspur habitat at times of year when it is considered most dangerous to cattle, exemplified by the toxic window concept [3, 4, 23]. While this approach has certainly helped many producers better understand larkspur toxicity dynamics, there is no evidence that it has reduced the overall number of deaths. There are many reasons for this, and interactions are complex and place-based, but we suggest that a reliance on a static view of palatability is largely to blame.

An alternative to fight or flight is to manage grazing such that larkspur intake remains below the threshold where there is an observable negative effect on the cattle. This study provides an indication that this may be possible even in pastures with dangerous amounts of Geyer’s larkspur. For the first time, this model suggests that herd cohesion and stocking density are key drivers of larkspur-induced toxicosis, and that management decisions that influence these factors hold potential to limit deaths. Of crucial importance is the observation that herd cohesion, which has received almost no consideration in the broader grazing management literature, is an important determinant of risk of death from larkspur.

An essential point for understanding how increased herd cohesion and stocking density reduced deaths is that Geyer’s larkspur grows most densely in relatively productive areas, which are thus desirable areas for foraging. Functionally, increased herd cohesion and stocking density lead to increased competition for forage, making it more difficult for any individual to monopolize a resource- and larkspur-rich area. Additionally, increased herd cohesion leads to less wandering among individuals, making it less likely an individual cow will wander into a dense larkspur patch alone. Evidence for the danger of wandering behavior is found in the disproportionate death rate of cows with the role of independent. Lastly, increased stocking density does appear to lead to dilution, but in the form of lowered maximum individual daily intake rather than lowered mean individual daily intake.

Mechanistically, decreased risk of lethal acute toxicosis occurred through: 1) a narrowed distribution of individual daily alkaloid intake, 2) lowered mean and narrowed distribution of outlier alkaloid intake days. Herd cohesion played a stronger role in narrowing the distribution of daily intake, stocking density was more influential in lowering the mean of outlier intake, and both played a relatively equal role in narrowing the distribution of outlier intake events. Strong evidence for the role of these as mechanisms is provided by the much lower AICc score for the model with the mechanisms than for the model with HCF and SD (1386.3 vs. 1686.3). This suggests that other management interventions that succeed in influencing these mechanisms would have similar success in reducing deaths.

When we recognize that even in the worst-case scenario lethal acute toxicosis is a rare event among thousands of grazing-days, it becomes clear why narrowing the distribution of individual intake and reducing outliers is so important. With a mean MSAL-tolerance of 4,000 mg, an average bad day in a herd with low herd cohesion and low stocking density would put an individual (especially one with lower tolerance or higher attraction to larkspur) in danger. Meanwhile, individuals grazing in a herd with high herd cohesion and at a high stocking density in the same pasture, even those with low tolerance, would need at least a few upper-end intake days in a row to risk death—an unlikely occurrence.

Note that we selected the bounds of herd cohesion and stocking density to align with what we believe to be realistically achievable by managers in the western US. While stocking density is easily understood, it may be worthwhile to describe how we think the various levels of herd-cohesion-factor (HCF) could be achieved (reference Fig 3). We think of HCF values of 1 and 4 as representative of most current extensive management, such that there is a small to moderate amount of herding behavior but in which animals are often spread out across a large area. The difference between these two might be accounted for by differences in breeding history, carnivore pressure, or genetic drift. To achieve an HCF of 7, we think cattle would need to be selected for strong herding instinct or be regularly, but not necessarily continually, herded. An HCF of 10 is comparable to many herds of wild ungulates and is achievable through the continual presence of a herder or a sustained effort at selecting for herding behavior.

There are two additional ways that a rapid increase in herd cohesion may be achieved. First, a drastic increase in stocking density (via increased animal-units or subdivided pastures) to a level that approaches “mob” grazing can forcibly increase cohesion. Second, the emerging technology of virtual fencing holds tremendous promise for achieving rapid changes in grazing behavior, including herd cohesion [61].

An unexpected emergent phenomenon occurred at HCF=1, in the form of distinct persistent subherds (see Fig 3a). These subherds are small groups of >20 but usually <35 cows that stick closely together for an entire inter-watering period, with some exchange of individuals or combining when two groups meet. This does not occur at higher levels of HCF. Cows in distinct persistent subherds traveled significantly greater distances, spent more time seeking to be closer to herdmates rather than maximizing forage intake, and grazed more heterogeneously (Figs 6a-c). Nevertheless, these cows reached 150 AUMs of forage consumption in 94.3% of the model run time of cows at lower herd cohesion levels, suggesting higher grazing efficiency (Fig 6d). We believe that these data are evidence of a scale-dependent behavioral discontinuity that may hold relevance to other grazing management challenges [62].

### Model parsimony and study limitations

Perhaps the most obvious omissions from the model are those behaviors that we determined to hold little to no relevance to larkspur consumption, at least in this pasture. These include response to slope, resting, and some inconsistently understood aspects of dominance behaviors. While there is nothing preventing them from being included, we decided that in this case these behaviors would introduce uncertainty while adding little realism to cattle-larkspur dynamics. The model also excludes plant regrowth. For Geyer’s larkspur, this is not an issue, as plants that are clipped or grazed during the bud stage exhibit very little regrowth [K. Jablonski, pers. obs.]. For other forage, we determined that regrowth in July in this semi-arid climate would not be substantial enough within a single grazing period to warrant inclusion.

The occurrence and measurement of death in the model might strike some as unrealistic. However, given that deaths in a herd would change herd behavior in unknown ways, and that the owner of the cattle would likely intervene once one death-event had occurred, we believe that counting the death and resetting the cow’s MSAL-level is the most accurate way to assess risk at different management levels.

Another potential limitation concerns the model used for alkaloid metabolism. While there has been some effort at the generation of such a model [e.g., 48], these efforts have been limited to highly controlled settings using hay and other stored feeds and periodic dosing with alkaloids. Additionally, little to nothing is known about the role of other forage in exacerbating or mitigating the effects of larkspur consumption. As such, we had no confidence that a continuous metabolic model would be more useful than the simple daily half-life model that we used.

Despite these limitations, we are confident that we have realistically modeled cattle-larkspur dynamics, that increased herd cohesion and stocking density lower the risk of lethal acute toxicosis, and that variations in mean and maximum daily alkaloid intake are the predominant mechanism for this reduction. However, the exact values for when risk approaches zero may be dependent on the circumstances of this model iteration—that of *D. geyeri*, at the input values for mass and toxicity, on a ranch in northern Colorado.

It is worth noting that dangerous levels of *D. geyeri* are typically found on a limited number of a single operation’s grazing units. This means that the inclusion of herding to increase herd cohesion, for example, would usually only be necessary for a relatively brief period. In addition, it means that any potential secondary effects of sub-lethal larkspur consumption, such as appetite suppression or lethargy (whether and how these would occur is unclear), would be of similarly limited duration. Nevertheless, in pastures with a dangerous amount of larkspur, negative sub-lethal effects may be unavoidable even (or especially) when death is avoided.

As with any research where cattle lives and producer livelihoods are at stake, it is most important to emphasize that producers should exercise caution when incorporating our findings into their own management, including careful assessment of other potential effects of increased herd cohesion or stocking density. Those with low amounts of Geyer’s larkspur or with no history of losses might find comfort in altering their grazing management to incorporate this study’s findings. Those with a great deal of larkspur (Geyer’s or other species) or a history of losses should be more careful.

### Other model implications and future directions

There is a broad literature on the effect of stocking rate/stocking density on many outcomes (though not larkspur-induced toxicosis) but very little on the effects of herd cohesion, nor on the interaction of these factors [44]. This is likely due to the relative ease of varying cattle numbers versus manipulating cattle behavior. Because this study provides evidence that it is not only the number of animals but also how they behave that affect the likelihood of death by larkspur, we are excited to explore the role of herd cohesion, particularly the emergent property of distinct persistent subherds, in other aspects of grazing ecology. If herd cohesion is genetically encoded, matrilineally-oriented, or management-determined (or a combination thereof), what role might it play in other negative outcomes, such as overgrazing of riparian areas or exposure to predation by carnivores [63], and how might we influence it in different scenarios? The evolving promise of affordable GPS tags means that we may also start to be able to test this through direct observation of entire herds [64].

For cattle-larkspur dynamics, our next step is to place these modeling results in context with ongoing plant experiments and producer surveys to better formulate management recommendations that work. Additionally, we would like to improve our understanding of alkaloid metabolism and tolerance, as well as the role of preference in larkspur intake. For alkaloid metabolism and tolerance, this means building upon previous studies [e.g., 21], which have been undertaken in highly controlled settings using periodic high dosing, to model the stochastic dosing in a dynamic environment that occurs in reality. For larkspur preference, this means moving beyond the entirely anecdotal evidence of bouts of larkspur consumption [e.g., 65] to a more sophisticated understanding of the role of preference, diet mixing, and satiation in larkspur-induced toxicosis [66, 67].

A final next step for the model presented here is what Augusiak et al. [19] term model output corroboration, wherein model outputs are compared to new, independent data and patterns. As noted above, this is very difficult when cattle lives are at risk. However, the results presented here have encourage us to start to think about how such corroborative data could be collected. This will likely entail a combination of full-herd GPS with careful on-the-ground monitoring by a herder.

Though ABMs have some limitations, we believe they offer an exciting new tool for understanding the grazing behavior of livestock. Indeed, the synergistic emergence of financially viable GPS technology [64] and “virtual fencing” [61], along with the increasing power of desktop computers, suggests that the time is right for a computational revolution in livestock grazing management. We are excited that this study provides a first example of the potential of agent-based models to contribute to this revolution.

## Acknowledgements

We thank Joel Vaad, manager of the Maxwell Ranch, for access, assistance, and advice. Early conversations with Michael A. Smith and the Sims family of McFadden, WY, provided crucial insight into the relationship between grazing management and larkspur. Tanner Marshall assisted in data collection and asked difficult questions that moved the work forward. We are grateful to DigitalGlobe for a generous imagery grant that greatly improved our ability to assess the grazing landscape, and to Hexagon Geospatial for providing the software to analyze the imagery.

